# Geometric eigenmode brain fingerprinting and its longitudinal associations with adolescent mental health and wellbeing

**DOI:** 10.1101/2024.08.08.607260

**Authors:** Amanda Boyes, Paul E. Schwenn, Zack Y. Shan, Mayuresh Korgaonkar, James Pang, Taliah Prince, Lia Mills, Michelle F. Kennedy, Daniel F. Hermens

## Abstract

**Background:** ‘Brain fingerprinting’ research posits that individual uniqueness can be identified by structural and functional features that may also be linked to mental health outcomes. Global structural features of the brain can be succinctly and directly captured from magnetic resonance imaging (MRI) via the eigenmodes of the cortical surface - known as *geometric eigenmodes*. This research investigates how the uniqueness of geometric eigenmodes changes across adolescence and their longitudinal relation to mental health and wellbeing.

**Methods:** The current study utilised n=613 MRI, self-report and demographic datasets from N=116 community-recruited adolescents enrolled in the Longitudinal Adolescent Brain Study (LABS), between the ages of 12-17 years. MPRAGE scans at each participant’s visit were used to derive 225 left-hemisphere geometric eigenmodes. Eigenmodes were clustered into 14 eigengroups and developmental trajectories of their uniqueness and longitudinal associations with mental wellbeing and psychological distress were examined.

**Results:** All eigengroups become significantly more unique longitudinally, and higher mode (shorter wavelength) eigengroups were more unique than lower mode groups in adolescence. Less uniqueness in ‘eigengroup 6’ was significantly associated with higher psychological distress and lower mental wellbeing at concurrent and future timepoints.

**Conclusion:** Geometric eigengroup brain fingerprinting offers a novel way to examine neurodevelopment. This study provides evidence that eigengroups have distinct trajectories from adolescence to adulthood, consistent with other imaging studies demonstrating increasing uniqueness in this period. Importantly, they are associated with mental health state and thus may represent neurobiological markers for mental illness onset, building on previous LABS research demonstrating that the functional uniqueness of the ‘cognitive control network’ predicts psychological distress four months later.

## 1. Introduction

‘Brain fingerprinting’, or the ability to determine what makes an individual’s brain unique in terms of structure or function, has been an important pursuit of neuroscientists for over a decade (Chen et al., 2022; Chen et al., 2024; Finn et al., 2015). Different methods have been utilised to understand how neurobiological uniqueness manifests, how much is heritable or influenced by environment, and how this is linked to behavioural or cognitive outcomes (Chen et al., 2022; Mueller et al., 2013). However, the optimal way of determining neurobiological uniqueness and how this may be linked to (or predict) mental health symptoms (and thus useful targets for personalised mental health care), remains under-researched (Hermens et al., 2023).

Functional connectomes, which map the brain’s functional connectivity (FC), can be used to identify similarity-to-self, similarity-to-others, and individual uniqueness in adolescents and adults (Finn et al., 2015; Shan et al., 2022b; St-Onge et al., 2023). Recently, Shan et al. (2022b) examined the development of functional connectome uniqueness, and how uniqueness among known FC networks was related to mental health in adolescents (Shan et al., 2022b). These analyses revealed those with a less-unique cingulo-opercular network (CON; also referred to the ‘cognitive control network’) had higher psychological distress four months later. These findings provide the first indication that maturational delays in the development of networks associated with executive functioning may predict subsequent increased mental health problems (Hermens, 2023; Hermens et al., 2023).

Recent research examining the stability of these uniqueness metrics over the lifespan suggests that while brain fingerprinting can reliably identify individuals in different age groups, self-similarity and similarity-to-others may develop in an inverted-U trajectory, with both metrics being higher in younger (<∼30 years) and older (>∼70 years) age groups (St-Onge et al., 2023). Further, functional connectome uniqueness may involve different features for different individuals and be influenced by neuropsychiatric symptoms or disorders (St-Onge et al., 2023). Geometric eigenmodes, which can be extracted from structural MRI scans, offer a promising alternative to traditional parcellation approaches, as the technique removes issues with variability and lack of comparability between approaches (Müller et al., 2022; Pang et al., 2023). Cortical shape has also been suggested to be akin to a brain fingerprint, with uniqueness in shape asymmetry between the left and right hemispheres at scales of ∼37mm able to optimally identify individual adults (Chen et al., 2022). More specifically, geometric eigenmodes - as a recent application of neural field theory - are utilised to understand structure-function relationships, as they match macro-scale structural cortical features with functional activity within the brain (Robinson et al., 2016).

Geometric eigenmodes represent fundamental patterns of spatial oscillations on the cortical surface (Robinson et al., 2016). The eigenvalues associated with these eigenmodes are related to their spatial characteristics, with larger eigenvalues generally corresponding to more rapidly and locally varying spatial patterns (Pang et al., 2023). The degenerate eigenvalues of a sphere can be used as a basis to group cortical eigenmodes with similar spatial patterns of oscillation (Cao et al., 2024; Robinson et al., 2016). Recent research by Cao et al. (2024) and Pang et al. (2023) utilised left hemisphere surface-based eigenmodes in adult samples to examine associations with other MRI measures of function and structure. Pang et al. (2023) found that: (i) uni-hemispheric geometric eigenmodes of the cortical surface are more useful than other, more complex connectome measures in understanding brain dynamics; and (ii) coarser wavelength modes (∼60mm) appear to be most important in accurately reconstructing a range of task-based and resting-state brain activity. Further, Cao et al. (2024) found that 225 left hemisphere eigenmodes, classified into 14 eigengroups, can be used to examine phenotypic morphometric differences between groups (e.g., by sex, or patient versus control), cross-sectionally, across different research samples, capturing important cortical shape-based information that is less affected by noise. Thus, structural analyses utilising geometric eigenmodes appear to yield more consistent results than usual metrics of brain surface structure (offering a useful alternative to traditional methods) (Cao et al., 2024). Eigenmodes may also be utilised to examine neurodevelopment and how disruptions in them relate to mental health disorders or symptoms (or vice-versa) (Pang et al., 2023).

Adolescence is characterised by rapid biological, social, emotional, and cognitive changes, presenting dynamically changing health and developmental needs. This period is recognised as crucial for brain development and mental health risk, as well as opportunity for intervention (see review: King, 2019). Altered brain development trajectories have been linked to the onset and development of psychiatric symptoms in adolescents (Bick & Nelson, 2016; Whittle et al., 2013). Global prevalence rates from 2019 indicate that 12.4% of young people aged 10-14 years and 13.96% of 15-19 year-olds had at least one mental disorder (Kieling et al., 2024). These rates may be higher when examined at the symptom-level, as a survey of mental health and wellbeing in Australia found that 40% of people aged 16-24 reported experiencing symptoms of a diagnosed disorder in the last 12 months, the highest rate of any decade of life (Australian Bureau of Statistics, 2022). Further, young Australians aged 16-34 were most likely (20%) to experience high or very high levels of psychological distress (Australian Bureau of Statistics, 2022), and research across 73 countries examining both mental wellbeing and psychological distress have found that on average, females adolescents have poorer mental health across measures (Campbell et al., 2021). Meanwhile, concurrent brain structural changes occur during adolescence, including generalised pruning of grey matter volume, a decline in mean cortical thickness, and increased myelination of white matter tracts (Gatt et al., 2014; Jamshidi et al., 2020).

Recent studies examining how changes in structural features may be associated transdiagnostically or in emerging disorders have tried to identify the most useful metrics and methods for future research that will enable the most helpful information for interventions (Hettwer et al., 2022; Van Rheenen et al., 2023). Research examining cortical surface area, volume, thickness, and gyrification as well as ‘brain age’ models have found different associations with mental health outcomes in adolescents and adults (Drobinin et al., 2022; Hettwer et al., 2022; Van Rheenen et al., 2023). Specifically, a study by Van Rheenen et al. (2023) examined cortical surface area, thickness and gyrification following first episode psychosis in *n*=35 participants aged 15-25 years, finding that increased surface area may be key to understanding psychosis risk in the developing brain. Research utilising cortical grey matter volume and surface area measurements to develop a ‘brain age’ model found that adolescents with major depressive disorder and worse functional outcomes (across six study cohorts aged between 9-19 years) had ‘older’ brains to their peers (Drobinin et al., 2022). While other research has tried to identify commonalities across multiple diagnoses, finding that there are shared effects of mental illness on cortical thickness in adults (Hettwer et al., 2022). Despite such evidence, there is not yet a consensus on the most effective way to utilise structural features of the brain to distinguish between those who may be at an increased risk for mental health problems from those who are on a developmental trajectory with consistent mental wellbeing.

An international consensus statement has highlighted the global need for accurate and useful transdiagnostic research to inform early interventions and support services, incorporating measures such as biological or other comorbid symptoms (Shah et al., 2020). To this end, recent research has emphasised the value of including measures of positive mental health and functioning, beyond indicators of negative mental health, for use in community prevention, biological research and clinical treatments (Boyes et al., 2022; Gatt et al., 2018; Park et al., 2022). Measures of wellbeing, such as quality of life scales, can provide important information about key areas of need in a young person’s life, as well as areas that may offer protective buffers against mental ill-health risks (Cotton et al., 2022). The COMPAS-W, a measure of eudaimonic and hedonic wellbeing, has been shown to correlate with other measures of mental health in early adolescence (Beaudequin et al., 2020; Lam et al., 2024) and has also been used to identify differences in subcortical grey matter volume in adolescents and adults (Boyes et al., 2022; Gatt et al., 2018; Park et al., 2022). The Kessler Psychological Distress Scale (K10) captures current experience of psychological distress, and is one of the most widely-used and well-validated screening tools for psychological symptoms and distress, utilised in a range of adolescent neuroimaging studies to date (Boyes et al., 2024; Iorfino et al., 2017; Jamieson et al., 2022; Shan et al., 2022b). In addition to the value of assessing both mental wellbeing and psychological distress, there is also great utility in conducting multiple assessments over time, to gain an accurate representation of an individual’s variability and trajectory in mental health (Welsh et al., 2020). This may be particularly useful in adolescence, which is characterised by substantial changes in mental health.

Geometric eigenmode uniqueness has not yet been examined in terms of development or associations with mental health in adolescence. However, some key related findings by recent research by Chen et al. (2022; 2024) provide guidance for future research, as they found that: (i) higher cortical asymmetry uniqueness is associated with positive and negative symptoms in young people aged 16-34 years with early psychosis, at coarse scales (wavelengths ∼40mm to ∼75mm), with eigengroups 6-9 observing the largest effect sizes; and (ii) in adults (22-35 years) uni-hemispheric shape is highly heritable at coarse scales (wavelength ∼65mm), however, this declines with higher mode groups (i.e., above eigengroup 6). Furthermore, Pang et al. (2023) demonstrated that left hemisphere geometric eigenmodes accurately represent functional brain activity. Such evidence, along with findings of reduced FC uniqueness predicting subsequent increased psychological distress [K10 scores] (Shan et al., 2022b), suggests that the assessment of geometric eigenmodes and their associations with mental health symptoms in adolescence is warranted.

Further, to aid in the untangling of research to date, it may be helpful to categorize eigengroups into ‘low’ spatial frequency (i.e., groups 1-5), ‘mid’ spatial frequency (groups 6-9) and ‘high’ spatial frequency (groups 10+). This is suggested as ‘low’ to ‘mid’ groups (i.e., the first ∼100 eigenmodes) may be most useful in understanding brain function, while ‘mid’ to ‘high’ groups may be most useful in understanding mental health symptoms and environmental influences, as heritability is lower in these higher modes (Chen et al., 2022; Chen et al., 2024; Pang et al., 2023). Accordingly, this longitudinal study analysed data collected across 15 timepoints, over a 5-year period, for adolescent participants enrolled in LABS between the ages of 12-17 years. The data included self-reported measures of wellbeing and psychological distress (COMPAS-W and K10), as well as left hemisphere geometric eigenmode data extracted from structural MRI scans at each timepoint. Drawing on the literature, we hypothesise that: (1) left hemispheric geometric eigengroups will become increasingly unique in adolescence; and (2) ‘mid’ to ‘high’ frequency geometric eigengroup uniqueness will be associated with mental health.

## 2. Method

### 2.1 Study Design and Participants

Ethics approval was granted by the UniSC Human Research Ethics Committee (Approval A181064). This study utilised self-report questionnaire and magnetic resonance imaging (MRI) data collected over the first five years of LABS between 18 July 2018 to 3 July 2023, design and recruitment details are described elsewhere (Beaudequin et al., 2020; Boyes et al., 2022; Levenstein et al., 2023). Written consent was obtained from all parents/caregivers and participants. Assessments took up to five hours to complete, with scheduled breaks, and were completed with trained researchers at the Thompson Institute, UniSC. A sample of *N*=116 young people attended a minimum of two (maximum of 13) timepoints (TPs) between baseline (TP1) and final (TP15) LABS visits, at 4-monthly intervals, resulting in a total of *n=*613 datasets.

Participants (58 female; 9 left-handed, 3 ambidextrous) were ∼12 years old at TP1 and 17 years old at TP15; see Table S1 Supplementary Materials for summary of demographics for data included across all timepoints.

### 2.2 Inclusion and Exclusion Criteria

Selection criteria included participants aged 12 years, in their first year of secondary school (Grade 7) and proficient in spoken and written English. Young people were excluded prior to entry into the study if they: suffered from a major neurological disorder, intellectual disability, or major medical illness; had sustained a head injury (with loss of consciousness more than 30 minutes); or if they were unable to complete the MRI component.

### 2.3 Measures

#### 2.3.1 Self-reported Wellbeing (COMPAS-W)

As part of the self-report questionnaire, participants completed the 26-item COMPAS-W scale, a reliable indicator of mental wellbeing (Gatt et al., 2014). Each question was answered on a 5-point Likert-type scale (1 = *strongly disagree* to 5 = *strongly agree*) resulting in a ‘total wellbeing’ score of 26-130, with higher scores indicating greater wellbeing. The COMPAS-W has been validated for use in 12-61 year-olds and includes three measures of eudaimonic (psychological) wellbeing (i.e. ‘own-worth’, ‘mastery’ and ‘achievement’), and three measures of hedonic (subjective) wellbeing (i.e. ‘composure’, ‘positivity’ and ‘satisfaction’) (Gatt et al., 2020; Gatt et al., 2014). Previous research has identified that individuals who experience ‘languishing’ wellbeing are clinically distinct, experiencing more depressive symptoms, poorer functional outcomes, and different genetic profile to those who are ‘moderate’ or ‘flourishing’ (Gatt et al., 2014; Jamshidi et al., 2020).

#### 2.3.2 Self-reported Psychological Distress (K10)

Participants completed the K10 scale, with 10 items answered on a 5-point scale (1 = *none of the time* to 5 = *all of the time*), and an overall score of 10-50, with higher scores indicating greater psychological distress. The K10 is a valid measure of psychological distress in adolescents over the previous 30 days, which uses the participant’s total score to identify their level of non-specific psychological distress and likelihood of psychological disorders, particularly depression and anxiety (Andrews & Slade, 2001; Chan & Fung, 2014; Kessler et al., 2003; Lawrence et al., 2015; Sunderland et al., 2011). The Australian Bureau of Statistics categorises K10 scores of 10-15 as ‘low’, 16-21 ‘moderate’, 22-29 ‘high’ and 30-50 ‘very high’(Andrews & Slade, 2001; Australian Bureau of Statistics, 2012).

#### 2.3.3 Magnetic Resonance Imaging

Participant MRI brain scans were acquired using a 3-Tesla Siemens Skyra scanner (Erlangen, Germany) with a 64-channel head and neck coil, performed at the Nola Thompson Centre for Advanced Imaging (Thompson Institute, UniSC). As part of the MRI protocol, 3D whole brain structural imaging was acquired using a T1-weighted magnetization prepared rapid acquisition gradient echo sequence (MPRAGE: TR=2200ms, TE=1.71ms, TI=850ms, flip angle=7°, spatial resolution=0.9x0.89x0.89mm, FOV=208x256x256, TA=3:57).

### 2.4 MPRAGE and Eigenmode Analyses

#### 2.4.1 Structural Processing

T1-weighted images were processed using the FreeSurfer (7.4.0) longitudinal recon-all pipeline to conduct CROSS (cross-sectional), BASE (within-subject), and LONG (longitudinal) analyses for each participant (Reuter et al., 2012). Software was packaged in singularityCE (version 3.8.0), on the UniSC HPC system, using PBS (version 20.0.0), Ubuntu 20.04, and connected to a CEPH cluster (version 15.2.13). MRI structural data was extracted from the CROSS and LONG brains. Euler numbers were calculated for all MRI scans using the below equation (1):

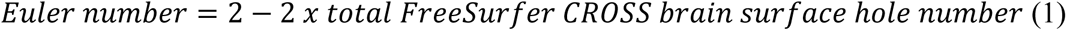

Euler numbers provide an indication of cortical surface topology and have been used in neurodevelopment studies as a measure of image quality (Dale et al., 1999; Rosen et al., 2018). A perfect surface with no holes would have an Euler number of two, however, in adolescent imaging studies, it has been previously reported that values can be as low as -380 and may mediate apparent relationships between grey matter and age (Rosen et al., 2018).

#### 2.4.2 Eigenmode Calculations and Processing

All eigenmode analyses were undertaken in purpose-built singularity containers on the UniSC HPC system. Geometric eigenmodes for the left cortical hemisphere were calculated using the method developed by Pang et al. (2023), whereby (i) midthickness surface files were created using the FreeSurfer-generated pial and white surfaces, converted to fs_LR_32k space (with 32,492 vertices) and vtk format; and (ii) these files were run through the surface eigenmode pipeline, using the provided github script, with no binarized medial wall mask, to extract 225 eigenmodes and corresponding eigenvalues through the eigen-decomposition of the Laplace-Beltrami operator (LBO), which is given by equation (2):

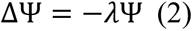

where Δ is the LBO and Ψ is an eigenmode with corresponding eigenvalue λ. Eigenmodes of the cortical surface describe the harmonic modes of oscillation, analogous to how the eigenmodes of a drumhead describe the harmonic modes of oscillation for a drum skin when struck with a stick. However eigenmodes here describe the patterns of neuronal activity rather than mechanical vibrations (Robinson et al., 2016). Pang et al. (2023) has shown that 200 eigenmodes are sufficient to accurately represent macroscale brain activity, while Cao et al. (2024) included a further 25 modes to create 14 eigengroups (Table S2). These groups are derived from the spherical version of the cortex (Robinson et al., 2016). Solving the eigenmodes of a sphere result in degeneracies, wherein certain eigenmodes have identical eigenvalues, creating an approximate basis for grouping cortical eigenmodes (see Supplementary Materials, equation 1). The 14 eigengroups were used to calculate longitudinal uniqueness for each scan, using a similar method to that outlined in Shan et al. (2022a), see Figure 1. This involved calculating (1) similarity-to-self and (2) similarity-to-others over time for each eigenmode matrix (with 225 eigenmodes x 32,492 vertices for each scan). These similarities were calculated using Python, see equations 3-5 below. Because higher order eigenvalues are closely spaced, there is the potential for ‘mode swapping’ where the eigenmodes for one individual may not match that of the other (Chen et al., 2024; Robinson et al., 2016). To overcome this, eigenmodes were matched by maximum Pearson correlation before calculating uniqueness.

**Figure 1:**
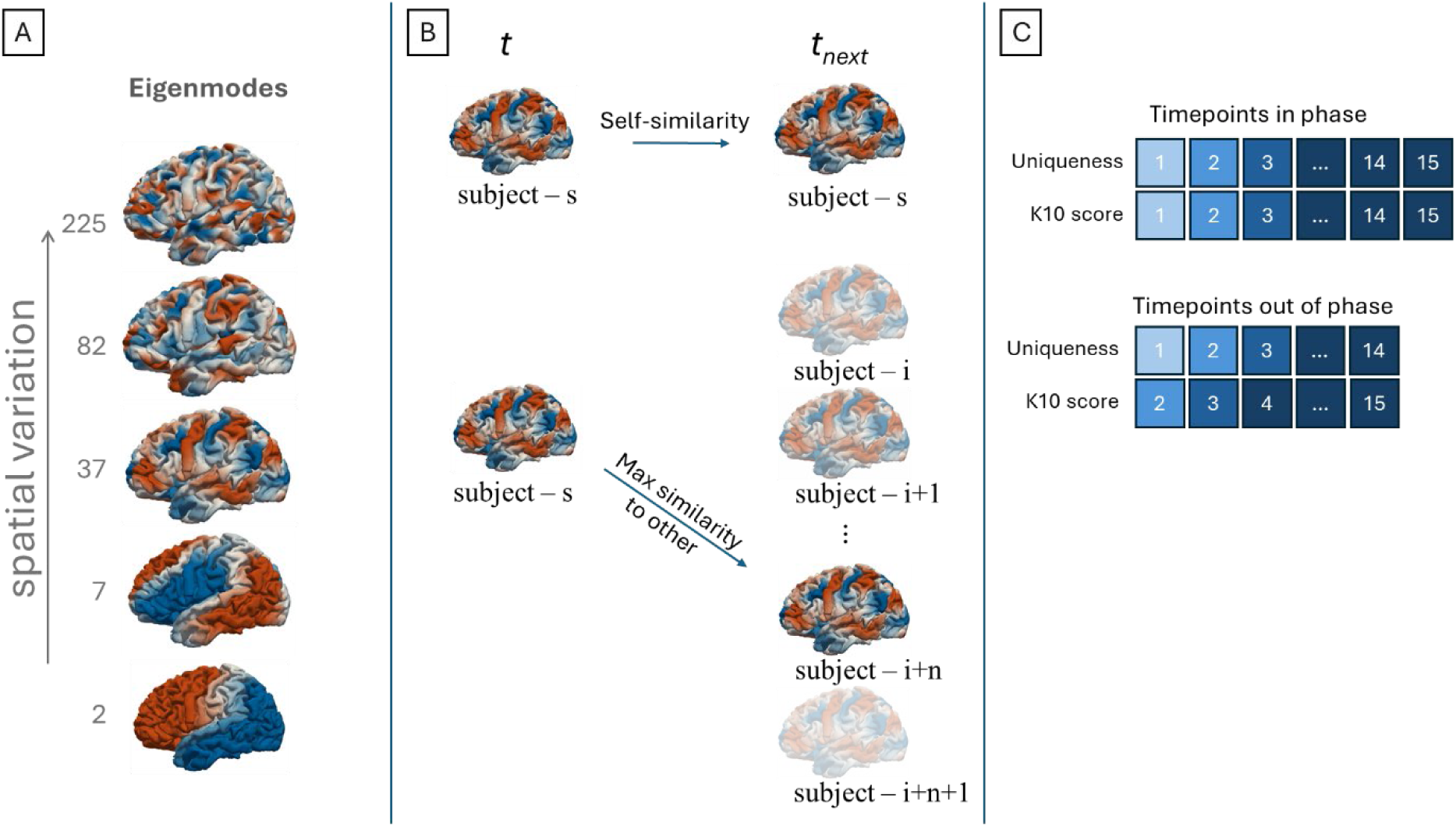
**A)** Representative surface eigenmodes showing increased spatial variation in higher order eigenmodes (white regions represent values around zero - nodal lines). **B)** Schematic showing similarity to self at the next timepoint (upper images) and maximum similarity to others (lower images), average of each pairwise correlation of eigenmodes within each eigengroup. **C)** Schematic showing in- and out-of-phase (lagged) timepoint alignment to associate uniqueness with concurrent (upper) and future (lower) mental health (e.g., K10 score).

##### Self-similarity

For each subject, self-similarity (i.e., the similarity between eigengroups from the same individual but different TPs) was calculated between consecutive available timepoints for each eigengroup using the absolute value of the Pearson correlation coefficient, see equation (3):

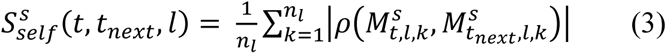

Where *s* is the subject index, *t* is the current timepoint, *t*_*next*_ represents the next available timepoint from timepoint *t* for subject *s* (accounting for potentially missing data), *l* is the eigengroup index, *k* is the eigenmode index in eigengroup *l*, *n*_*l*_ is the number of eigenmodes in the eigengroup, 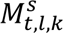 is the *k*-th eigenmode of eigengroup *l* at time point *t*, for subject *s* and ρ(⋅,⋅) is the Pearson correlation coefficient.

##### Similarity-to-others

Similarity-to-others (i.e., the similarity between an eigengroup of a given individual and eigengroup from a different subject and TP) was computed by comparing each subject’s eigenmode matrix at one time point to the maximum of all other subjects’ matrices at the subsequent time point (the same time point as the subjects next time point), see equation (4):

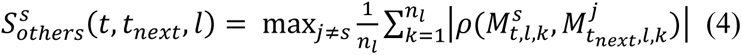

where *j* indexes over all other subjects except the current subject *s*.

##### Uniqueness

Uniqueness was then calculated as the ratio of self-similarity to the similarity-to-others, see equation (5) quantifying how distinct a subject’s eigenmode patterns are compared to others over time. Thus, uniqueness values greater than 1 indicate that eigenmodes are more similar within a given subject over time, and therefore identifiable (increasing uniqueness).

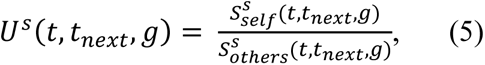

### 2.5 Statistical Analyses

#### 2.5.1 Data Screening

All outcome variables were confirmed to comprise valid scores so there were no artefactual outliers. Data quality assessment for the MRI scans involved exclusion due to prior- identified artefact (e.g., braces or excessive motion), resulting in an initial sample of *n*=622 scans. Following FreeSurfer analysis, the most extreme Euler numbers (z scores <- 3, indicating poorest quality scans) were used to identify additional scans that should be visually inspected. Raw scans were then used to determine whether data quality was acceptable and could retained (i.e., did not contain braces or motion artefact); or excluded prior to analyses. Following removal of the *n*=9 poorest-quality scans, a final check of the *n*=613 scans included in the analyses found that *n*=6 had Euler number z scores <-3 (raw Euler values -172, -174, -186, -198, -202, -236), however, these were retained as their Euler numbers were still within acceptable ranges, and visual inspection of the raw scans supported their inclusion. The final sample included Euler numbers ranging from -236 to -8 (mean -71.77, SD 31.95). Statistical analyses and plots were carried out using SPSS Statistics for Windows, Version 29.0.0 (IBM Corp., Armonk, NY) and R Studio version 4.2.

#### 2.5.2 Generalised Estimating Equations Analyses

Longitudinal development of eigengroup uniqueness and associations with wellbeing and psychological distress were determined using Generalised Estimating Equations (GEE). Prior to analysis, to aid with interpretation, age was centred to 12 years. In line with previous analyses (Boyes et al., 2024; Shan et al., 2022a, 2022b) a full GEE model was conducted with the following settings: the participant ID_TP as the subject variable; exchangeable and AR(1) working correlation matrices (to allow selection of best model fit for interpretation); a linear link function and chi-square statistics for model effect testing. Models examining longitudinal uniqueness development included uniqueness for eigengroups 1-14 as the dependent variable; sex as the factor; age (centred at 12 years) and Euler numbers as covariates. Models examining associations with mental health and wellbeing included K10 or COMPAS-W total scores (at current or next future timepoint, i.e., lagged analyses) as the dependent variable; eigengroups 6- 14 uniqueness, age (centred at 12 years) and Euler numbers as covariates; and sex as the factor. Euler numbers were used as a covariate to confirm whether any significant associations remained after controlling for MRI data quality, and the inclusion of only eigengroups 6-14 uniqueness in the models with mental health reduced the number of parameters, increasing power, and was supported by previous research indicting these ‘mid’ to ‘high’ spatial frequency eigengroups are most important to examine.

## 3. Results

### 3.1 Eigenmode Uniqueness

As uniqueness requires two timepoints (i.e., the current and next timepoint for an individual, or the current timepoint for an individual and next timepoint for others), there were *n*=497 uniqueness values calculated for the *N*=116 participants for each of the 14 eigengroups (minimum 1-maximum 12 uniqueness values per participant). Table S3 shows that higher eigengroups (i.e., those with shorter wavelengths) had higher uniqueness values. Most uniqueness values (77.46%, *n*=385) were calculated between adjacent timepoints (∼4 months apart), however, the largest difference between timepoints was ∼44 months (i.e., 3.67 years, for *n*=1). There were no uniqueness values calculated for TP15, as each uniqueness value requires a future timepoint as a comparison.

#### 3.1.1 Developmental trajectories

Exchangeable and AR(1) GEE models had identical QICC (best fit) and results. Results (Table S4) revealed that all eigengroups become significantly more unique as adolescents aged (controlling for Euler number and sex). Eigengroups 1-8 were significant at *p <*.001 but with small effect sizes (*β* = 0.0 to 0.02); eigengroups 10, 11, 12 and 14 were significant at *p* ≤ .01 (*β* = 0.02 to 0.04); and eigengroups 9 and 13 were significant at *p* < .05 (*β* = 0.01 and 0.03 respectively). Uniqueness for the ‘low’ spatial frequency eigengroups (1-5) showed less variation, and further, these groups were less unique over time (i.e., uniqueness remained closer to ∼1) (Table S3). ‘Mid’ (6-9) and ‘high’ (10-14) spatial frequency eigengroups had higher uniqueness values and increased variability amongst individuals. Females had slightly higher uniqueness in eigengroups 1 and 10 at *p* < .05 (*β* = 0.001 and .04 respectively), and eigengroup 2 at *p* < .01 (*β* = 0.001).

#### 3.1.2 Associations between uniqueness and mental health

GEEs with AR(1) and exchangeable correlation structures were identical, results are reported in Tables S5-S8. Figure 2 highlights the key findings between eigengroup uniqueness and mental health at concurrent and future timepoints. Adjusting for age, sex and Euler number, less uniqueness in ‘mid’ spatial frequency eigengroup 6 was associated with both higher psychological distress (concurrent *β* = -14.8, *p* < .01; future *β* = -13.9, *p* < .01) and lower mental wellbeing (concurrent *β* = 30.7, *p* < .001; future *β* = 30.7, *p* < .01), with this relationship consistent for males and females. Three outliers for eigengroup uniqueness were identified (see males, 12- 14.5 years, Figure 2). These individuals were removed, and analyses rerun, with all results retained. Wellbeing significantly decreased (concurrent *β* = -1.6, *p* < .001) and psychological distress significantly increased (concurrent *β* = 1.1, *p* < .001) as participants aged, and there were significant sex differences in mental health. Females had *both* higher mental wellbeing (*β* = 2.34, *p* < .05) and psychological distress compared to males over time in the concurrent models (*β* = 1.35, *p* < .05) (Tables S5-S6). Lagged GEE results in Tables S7-S8 indicate similar patterns of mental health over time, however there was no longer a significant sex difference for mental wellbeing. Mental wellbeing and psychological distress for males and females over TP1-TP15 are summarised in Table S9.

**Figure 2:**
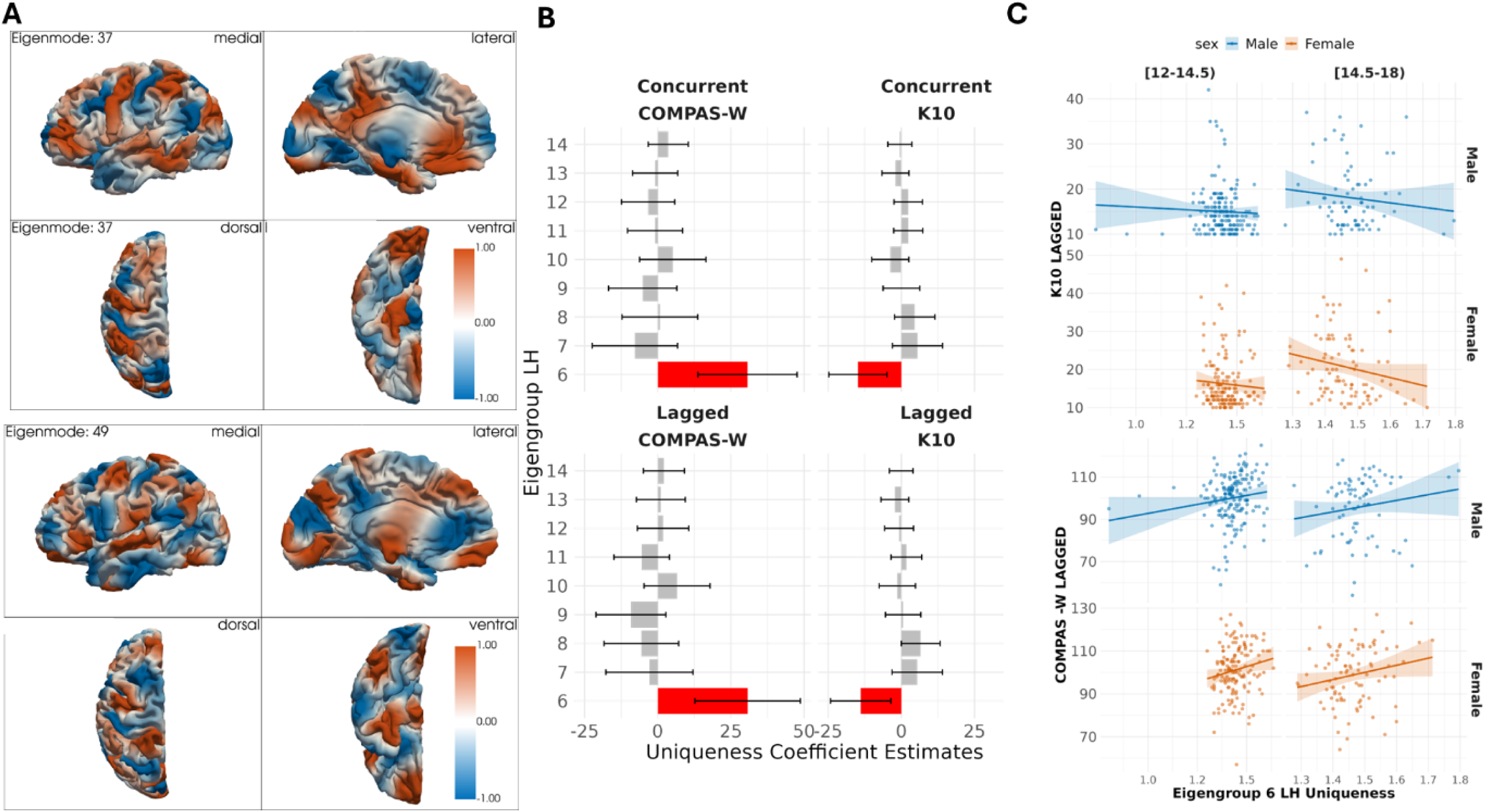
**A)** Representative eigenmodes in eigengroup 6: (top) eigenmode 37, (bottom) eigenmode 49, showing approximate length scales captured by this eigengroup (i.e., modes 37 – 49). **B)** GEE coefficient estimates for COMPAS-W (left) and K10 (right) with uniqueness for eigengroups 6-14: (top) concurrent and (bottom) out of phase (i.e., lagged). Coefficients are in the natural units of the measure. Eigengroup 6 was significant (red) while the others were not (grey). **C)** Scatterplots and trend line with 95% confidence intervals for K10 lagged (upper) and COMPAS-W (lower) by age 12-14.5 (left) and 14.5-18 (right) and males (upper, blue) and females (lower, orange) showing opposite trends in eigengroup 6 uniqueness with positive (COMPAS-W, wellbeing) and negative (K10, psychological distress) mental health measures.

## 4. Discussion

To our knowledge, this is the first time that left hemisphere cortical eigengroup uniqueness has been observed and demonstrated in adolescence, and then utilised as a predictor of current and future mental health symptoms and mental wellbeing, in a community cohort sample. The dataset was temporally rich, with a total of 613 MRI scans among *N*=116 adolescents, over a 5-year period (12-17 years old, with scan intervals as short as four months). Thus, the study is novel and provides a basis for a range of future research investigations.

Geometric eigenmodes constrain neuronal activity and provide benefits over other, more complex connectome measures in understanding brain dynamics, as they are more simple to obtain due to only requiring a structural mesh (Pang et al., 2023). Thus, understanding how the uniqueness of these eigengroups develops, and their associations with mental health over time, is a useful endeavour. As eigenmodes are linked to brain function, it was anticipated, based on previous functional connectome fingerprinting research (St-Onge et al., 2023), that geometric eigengroups would be unique in adolescence. This was confirmed, with shorter wavelength eigengroups being more unique than longer wavelength eigengroups, even when controlling for possible mode-swapping within eigengroups. The current findings also supported previous research by our team that brain fingerprints (via functional connectomes) are developing in terms of uniqueness during adolescence (Shan et al., 2022b). All 14 eigengroups became significantly more unique over time, albeit, with small effects sizes, and further, eigengroups 1-3 had effect sizes of ∼0 and constrained variability in uniqueness.

Eigengroups 1, 2 and 10 had significant sex differences over time, with females having higher uniqueness than males. However, eigengroups 1 and 2 had effect sizes of .001 and Table S3 indicates a lack of variance in uniqueness, thus it is unclear whether these findings can be utilised in a meaningful way. Chen et al. (2024) note that asymmetry uniqueness in eigengroup 1 provides little individual-specific information, and these groups were excluded from the analyses with mental health, thus, future research should determine whether uniqueness in these ‘low’ spatial frequency modes adds to our understanding of neurodevelopment (or not). Eigengroup 10 had increased variability in uniqueness, and the significant difference by sex had an effect size of .04 indicating that there is a small (but higher) possibility that subsets of ‘high’ spatial frequency mode groups may develop differentially by sex over time in adolescence.

Analyses of ‘mid’ to ‘high’ spatial frequency eigengroups 6-14 with mental wellbeing and psychological distress over time found that lower uniqueness in eigengroup 6 (including eigenmodes 37-49) was significantly associated with increased psychological distress and decreased mental wellbeing (at the same and future timepoints). Further, mental health and wellbeing significantly decreased over time in all GEE models, while females were significantly higher on both measures compared to males for concurrent data analyses. Previous research has highlighted the gap in mental health between males and females, in both psychological distress and wellbeing (Campbell et al., 2021), with different strengths and weaknesses identified by sex, indicating potential for specific targets for interventions (Matud et al., 2023). Examining fluctuations over time, males had higher psychological distress than females at timepoints 6 and 14, where they also had lower wellbeing than females (Table S9). This may indicate that there are different developmental, social or mental health related changes occurring from ∼14 years onwards that are worth examining in future studies. On average, males experienced more timepoints of poor mental wellbeing (i.e., at 12/15 timepoints), while females had higher average psychological distress at 13/15 timepoints. Mental health and wellbeing measures could be combined to create different profiles based on distress and wellbeing experience, and the associations with eigengroup changes in adolescence (Driver et al., 2024). Other factors, such as environmental (e.g., exposure to pollution), genetic (e.g., family history of mental illness) and broader demographic factors (e.g., socio-economic status) may also be important in identifying different associations between age, sex, eigengroups and mental health in adolescence (Yoon et al., 2023). Thus, future research should examine different sub-types of wellbeing (eudaimonic and hedonic) or profiles combining these measures (e.g., languishing but not distressed versus flourishing but distressed) to further interrogate any potential relationship with positive mental health.

The findings observed in this study are supportive of previous LABS findings, which found that less unique CON (cognitive control network FC) was associated with higher psychological distress 4 months later (Shan et al., 2022b), and indicates that maturation of eigenmodes with a wavelength ∼68 mm may be linked to current and future mental health in adolescence. ‘Low’ to ‘mid’ spatial frequency eigengroups (modes 1-50; eigengroups 1-6, with a wavelength of ∼60mm or longer) have been found to be useful in studies utilising left hemisphere cortical eigenmodes to reconstruct a range of functional brain activity in adults (Pang et al., 2023). The upper end of this cluster of modes (specifically, eigengroup 6) overlaps with the ‘mid’ to ‘high’ spatial frequency eigengroups that have been found to be most useful in examining associations between early psychosis symptoms and eigenmode asymmetry uniqueness in adolescents and adults aged 16-34 (Chen et al., 2024). Therefore, the current study provides further support for the importance of eigenmodes 37-49 and suggest possible interactions between cognition, mental health and neurodevelopment that warrant further analysis, particularly in adolescence, as this may provide contextual clues for research in adult cohorts and in those with emerging mental disorders.

The findings presented here support the inclusion of uni-hemispheric cortical eigengroup uniqueness in studies of adolescent neurodevelopment and mental health. By combining existing knowledge and processes, in a novel dataset, the results demonstrate the applicability of existing methods, and highlight opportunities for future research. Such future research could include: (1) examination of right hemisphere eigenmode uniqueness development and associations with mental health; (2) exploring hemispheric asymmetry uniqueness (Chen et al., 2024); (3) creation of groups to compare uniqueness and symptoms between different clinical or transdiagnostic cohorts (Cao et al., 2024; Chen et al., 2024); (4) using the sliding window approach to identify whether eigenmodes are associated with other modes outside their assigned groups and/or whether the eigengroups are the best fit for the adolescent data (Chen et al., 2024); (5) exploring alternative uniqueness metrics such as Weighted Spectral Distance for measuring shape dissimilarity (Konukoglu et al., 2013); (6) examining whether eigengroup uniqueness ‘peaks’ at a certain number of eigenmodes (i.e., identify the optimal number of eigenmodes/eigengroups to reliably determine an individual in adolescence (Chen et al., 2022)); (7) identify if specific brain regions or networks can be represented by the eigenmodes in eigengroup 6; (8) see if more specific models of eigengroup uniqueness development can be established and utilised to predict age (akin to a ‘brain age’ model); and (9) development of models incorporating other environmental, psychological and biological measures to increase our understanding of the non-heritable portion of uni-hemispheric eigengroup uniqueness. Research in other adolescent cohorts should aim to replicate the findings presented here, to add to our understanding of the development of cortical structural features in adolescence (Tamnes et al., 2017; Vijayakumar et al., 2016). As LABS continues to collect data from more adolescents in the older age range of our sample (i.e., 15-17 years) we may be able to shed further light on the developmental changes associated with cortical eigenmode uniqueness.

### 4.2 Conclusion

The current study found that a community cohort of adolescents aged 12-17 years had unique geometric eigengroups in the left hemisphere, across 14 eigengroups, and that decreased uniqueness of ‘mid’ spatial frequency eigengroup 6 was associated with higher psychological distress and lower wellbeing longitudinally. To our knowledge, this is the first time that these associations have been investigated, and thus are novel findings. Decreases in wellbeing and increases in psychological distress over time were observed at short (4-monthly) intervals, with females exhibiting both higher wellbeing and psychological distress compared to males.

Increased uniqueness, longitudinally, was observed in all eigengroups. Findings suggest that maturation of eigengroup 6 (modes 37-49, wavelength ∼68mm) is not only related to current mental health, but also predicts future mental health in adolescence. This cluster of ‘mid’ spatial frequency eigengroups overlaps with research identifying important modes for (i) reconstruction of brain activity across a range of functions in adults and (ii) asymmetric uniqueness and early psychosis symptoms; suggesting that understanding how these eigenmodes develop may be useful in identifying targets for early intervention. When considered in the context of existing research, this study highlights the benefits of studying both mental distress and wellbeing and with cortical structural development in adolescence. The use of geometric eigenmodes in addition to other cortical features has the benefit of capturing information about brain activity that can be mapped onto functional MRI data or utilised separately to identify potential targets for early mental health (or other) interventions.

## Acknowledgments

Thank you to the young people who gave their time to participate in this research. Thank you also to the Thompson Institute radiographers, LABS research assistants Marcella Parker, Arline Rathjen-Duffton and Shae Rendall, as well as the students and other staff who assisted with data collection. Pete Embleton and Simon Boyes were instrumental in successfully processing the MRI data on the UniSC HPC.

## Statements and Declarations

### Data availability

The datasets generated and analysed during the current study are not publicly available due the fact that they constitute an excerpt of research in progress but are available from the corresponding author on reasonable request.

### Code availability

Code to reproduce the analysis results and figures of this study will be openly available at https://github.com/thompsoninstitute/ upon publication of the article.

### Ethics approval

This study was performed in line with the principles of the Declaration of Helsinki. Ethics approval was granted by the UniSC Human Research Ethics Committee (Date: 9 May 2018/ Approval No.: A181064).

### Consent to participate

Written consent was obtained from all parents/caregivers and participants prior to participation.

### Consent to publish

All participants were informed of the research scope and inclusion of data to publish and could opt out at any time.

