## Supplemental Tables for "Geometric eigenmode brain fingerprinting and its longitudinal associations with adolescent mental health and wellbeing"

**Supplementary Materials**

**Table S1: Key Demographics by Timepoint (N=116, n=613 datasets)**

| Timepoint | Mean age in years (SD) | Male, *n* | Female, *n* | *N* | *K10 data* | *COMPAS-W data* | *MRI data* |
|---|---|---|---|---|---|---|---|
| 1 | 12.60 (0.31) | 44 | 37 | 81 | *80** | 81 | 81 |
| 2 | 12.92 (0.32) | 42 | 35 | 77 | 77 | 77 | 77 |
| 3 | 13.22 (0.32) | 32 | 30 | 62 | 62 | 62 | 62 |
| 4 | 13.62 (0.31) | 33 | 28 | 61 | 61 | 61 | 61 |
| 5 | 13.96 (0.32) | 25 | 28 | 53 | 53 | 53 | 53 |
| 6 | 14.22 (0.27) | 14 | 14 | 28 | *27** | 28 | 28 |
| 7 | 14.63 (0.28) | 26 | 19 | 45 | 45 | 45 | 45 |
| 8 | 15.00 (0.24) | 20 | 19 | 39 | *37** | *37** | 39 |
| 9 | 15.32 (0.29) | 18 | 14 | 32 | *31** | 32 | 32 |
| 10 | 15.65 (0.27) | 11 | 13 | 24 | 24 | 24 | 24 |
| 11 | 15.96 (0.25) | 12 | 15 | 27 | 27 | 27 | 27 |
| 12 | 16.25 (0.25) | 11 | 14 | 25 | 25 | 25 | 25 |
| 13 | 16.68 (0.27) | 11 | 13 | 24 | 24 | 24 | 24 |
| 14 | 17.04 (0.25) | 8 | 13 | 21 | *20** | *20** | 21 |
| 15 | 17.42 (0.24) | 4 | 10 | 14 | 14 | 14 | 14 |
| Total | - | 311 | 302 | 613 | 607 | 610 | 613 |

**Missing self-report data (i.e., K10, COMPAS-W or both, as indicated) for this timepoint, excluded from analyses examining mental health or wellbeing and MRI associations. All n=613 datasets were included in analyses that included MRI data only.*

**Table S2: Wavelengths by eigengroup**

| Eigengroup | Wavelength Mean (SD) | Eigenmodes included in eigengroup |
|---|---|---|
| 0 | – | 1 |
| 1 | 313.22 (9.97) | 2–4 |
| 2 | 180.84 (5.76) | 5–9 |
| 3 | 127.87 (4.07) | 10–16 |
| 4 | 99.05 (3.15) | 17–25 |
| 5 | 80.87 (2.57) | 26–36 |
| 6 | 68.35 (2.18) | 37–49 |
| 7 | 59.19 (1.88) | 50–64 |
| 8 | 52.20 (1.66) | 65–81 |
| 9 | 46.69 (1.49) | 82–100 |
| 10 | 42.23 (1.34) | 101–121 |
| 11 | 38.55 (1.23) | 122–144 |
| 12 | 35.46 (1.13) | 145–169 |
| 13 | 32.83 (1.05) | 170–196 |
| 14 | 30.57 (0.97) | 197–225 |

**Note.* Average wavelengths by eigengroup for the current sample were based on $Rs_t$, calculated from the estimated total intracranial volume from the cross-sectionally processed scan at each participant's timepoint (t) ($Rs_t$ mean 70.50mm; min. 59.05mm - max. 77.21mm).

The theoretical wavelength of each eigengroup was calculated in R Studio version 4.2, using the following equation (1):

$$wavelength\ (l, Rs_t) = \frac{2\pi Rs_t}{\sqrt{l(l+1)}} \qquad (1)$$

Where $l$ is the eigengroup number (each group comprises $2l + 1$ eigenmodes) and $Rs_t$ is the radius of a sphere calculated from each participant's eTIV at timepoint (t).

**Table S3: Left hemisphere eigengroup uniqueness for *N*=116, *n*=497 datasets**

| Eigengroup | Minimum | Maximum | Mean | SD | Wald 95% CIs |
|---|---|---|---|---|---|
| 1 | 0.99 | 1.06 | 1.00 | 0.00 | 1.00-1.01 |
| 2 | 0.98 | 1.04 | 1.02 | 0.01 | 1.01-1.03 |
| 3 | 0.90 | 1.17 | 1.07 | 0.02 | 1.02-1.11 |
| 4 | 0.88 | 1.37 | 1.16 | 0.04 | 1.08-1.25 |
| 5 | 0.78 | 1.57 | 1.28 | 0.06 | 1.16-1.40 |
| 6 | 0.80 | 1.80 | 1.45 | 0.08 | 1.28-1.62 |
| 7 | 0.84 | 2.00 | 1.68 | 0.11 | 1.46-1.89 |
| 8 | 0.85 | 2.27 | 1.91 | 0.14 | 1.63-2.18 |
| 9 | 0.88 | 2.52 | 2.15 | 0.16 | 1.83-2.47 |
| 10 | 0.85 | 2.87 | 2.34 | 0.19 | 1.97-2.72 |
| 11 | 0.97 | 3.00 | 2.55 | 0.23 | 2.09-3.01 |
| 12 | 0.92 | 3.44 | 2.74 | 0.26 | 2.22-3.26 |
| 13 | 0.92 | 3.57 | 2.91 | 0.31 | 2.29-3.53 |
| 14 | 0.95 | 3.79 | 3.04 | 0.34 | 2.36-3.72 |

**Table S4: GEE results for eigengroup uniqueness over time for *N*=116, *n*=497 datasets**

| Dependent Variable | B (Age) | 95% CI | | Hypothesis Test | |
|---|---|---|---|---|---|
| | | Lower | Upper | Wald Chi-Square | Sig. |
| Uniqueness Eigengroup 1 | 0.00 | 0.00 | 0.00 | 16.10 | <.001[ac]*** |
| Uniqueness Eigengroup 2 | 0.00 | 0.00 | 0.00 | 66.08 | <.001[bd]*** |
| Uniqueness Eigengroup 3 | 0.00 | 0.00 | 0.01 | 23.03 | <.001*** |
| Uniqueness Eigengroup 4 | 0.01 | 0.01 | 0.01 | 45.95 | <.001*** |
| Uniqueness Eigengroup 5 | 0.01 | 0.01 | 0.02 | 21.86 | <.001[a]*** |
| Uniqueness Eigengroup 6 | 0.02 | 0.01 | 0.02 | 24.77 | <.001*** |
| Uniqueness Eigengroup 7 | 0.02 | 0.01 | 0.03 | 20.27 | <.001*** |
| Uniqueness Eigengroup 8 | 0.02 | 0.01 | 0.03 | 14.26 | <.001*** |
| Uniqueness Eigengroup 9 | 0.01 | 0.00 | 0.02 | 4.05 | 0.04* |
| Uniqueness Eigengroup 10 | 0.02 | 0.01 | 0.03 | 9.36 | 0.00[c]** |
| Uniqueness Eigengroup 11 | 0.02 | 0.01 | 0.04 | 8.75 | 0.003** |
| Uniqueness Eigengroup 12 | 0.03 | 0.01 | 0.04 | 7.18 | 0.01** |
| Uniqueness Eigengroup 13 | 0.03 | 0.00 | 0.05 | 5.11 | 0.02* |
| Uniqueness Eigengroup 14 | 0.04 | 0.01 | 0.06 | 9.50 | 0.002** |

*Note.* B values are for age (centred at 12 years) for all AR(1) models with eigengroup uniqueness as the dependent variable. The intercept was significant at $p < .001$ for all models. [a]Euler number was significant at $p < .05$. [b]Euler number was significant at $p < .001$. [c]Females had higher uniqueness at $p < .05$. [d]Females had higher uniqueness at $p < .01$. *$p <.05$, **$p \leq .01$, ***$p \leq .001$

**Table S5: Concurrent K10 GEE by eigengroup uniqueness, age, sex for *N*=116, *n*=497 datasets**

| Parameter | B | 95% CI | | Hypothesis Test | |
|---|---|---|---|---|---|
| | | Lower | Upper | Wald Chi-Square | Sig. |
| (Intercept) | 21.25 | 10.46 | 32.03 | 14.90 | <.001*** |
| [sex=Female] | 1.35 | 0.17 | 2.53 | 5.00 | 0.03* |
| Uniqueness Eigengroup 6 | -14.84 | -24.61 | -5.06 | 8.84 | 0.003** |
| Uniqueness Eigengroup 7 | 5.52 | -3.02 | 14.06 | 1.60 | 0.21 |
| Uniqueness Eigengroup 8 | 4.58 | -2.23 | 11.39 | 1.74 | 0.19 |
| Uniqueness Eigengroup 9 | 0.04 | -6.30 | 6.37 | 0.00 | 0.99 |
| Uniqueness Eigengroup 10 | -3.78 | -10.17 | 2.61 | 1.34 | 0.25 |
| Uniqueness Eigengroup 11 | 2.38 | -2.65 | 7.40 | 0.86 | 0.35 |
| Uniqueness Eigengroup 12 | 2.37 | -2.47 | 7.21 | 0.92 | 0.34 |
| Uniqueness Eigengroup 13 | -2.01 | -6.59 | 2.58 | 0.74 | 0.39 |
| Uniqueness Eigengroup 14 | -0.50 | -4.50 | 3.50 | 0.06 | 0.81 |
| Euler Number | 0.01 | -0.01 | 0.02 | 0.46 | 0.50 |
| Age, centred at 12 years | 1.13 | 0.65 | 1.60 | 21.64 | <.001*** |

$*p<.05$, $**p\leq.01$, $***p\leq.001$

**Table S6: Concurrent Wellbeing GEE by eigengroup uniqueness, age, sex, for *N*=116, *n*=497 datasets**

| Parameter | B | 95% CI | | Hypothesis Test | |
|---|---|---|---|---|---|
| | | Lower | Upper | Wald Chi-Square | Sig. |
| (Intercept) | 71.72 | 52.63 | 90.82 | 54.21 | <.001*** |
| [sex=Female] | 2.34 | 0.20 | 4.48 | 4.57 | 0.03* |
| Uniqueness Eigengroup 6 | 30.67 | 13.76 | 47.59 | 12.63 | <.001*** |
| Uniqueness Eigengroup 7 | -7.83 | -22.47 | 6.81 | 1.10 | 0.29 |
| Uniqueness Eigengroup 8 | 0.72 | -12.21 | 13.64 | 0.01 | 0.91 |
| Uniqueness Eigengroup 9 | -5.16 | -16.86 | 6.53 | 0.75 | 0.39 |
| Uniqueness Eigengroup 10 | 5.13 | -6.28 | 16.53 | 0.78 | 0.38 |
| Uniqueness Eigengroup 11 | -0.93 | -10.34 | 8.47 | 0.04 | 0.85 |
| Uniqueness Eigengroup 12 | -3.32 | -12.43 | 5.80 | 0.51 | 0.48 |
| Uniqueness Eigengroup 13 | -0.91 | -8.63 | 6.80 | 0.05 | 0.82 |
| Uniqueness Eigengroup 14 | 3.58 | -3.28 | 10.43 | 1.05 | 0.31 |
| Euler Number | 0.00 | -0.04 | 0.03 | 0.01 | 0.93 |
| Age, centred at 12 years | -1.55 | -2.38 | -0.72 | 13.47 | <.001*** |

$^{*}p < .05$, $^{**}p \leq .01$, $^{***}p \leq .001$

**Table S7: Lagged K10 GEE by eigengroup uniqueness, age, sex for *N*=116, *n*=497 datasets**

| Parameter | B | 95% CI | | Hypothesis Test | |
|---|---|---|---|---|---|
| | | Lower | Upper | Wald Chi-Square | Sig. |
| (Intercept) | 19.06 | 8.45 | 29.66 | 12.41 | <.001*** |
| [sex=Female] | 1.40 | 0.21 | 2.59 | 5.32 | 0.02* |
| Uniqueness Eigengroup 6 | -13.90 | -24.06 | -3.74 | 7.19 | 0.01** |
| Uniqueness Eigengroup 7 | 5.44 | -3.11 | 13.99 | 1.55 | 0.21 |
| Uniqueness Eigengroup 8 | 6.61 | 0.06 | 13.16 | 3.91 | 0.05 |
| Uniqueness Eigengroup 9 | 0.62 | -5.50 | 6.73 | 0.04 | 0.84 |
| Uniqueness Eigengroup 10 | -1.37 | -7.59 | 4.85 | 0.19 | 0.67 |
| Uniqueness Eigengroup 11 | 1.73 | -3.56 | 7.02 | 0.41 | 0.52 |
| Uniqueness Eigengroup 12 | -0.76 | -5.68 | 4.16 | 0.09 | 0.76 |
| Uniqueness Eigengroup 13 | -2.19 | -6.91 | 2.53 | 0.83 | 0.36 |
| Uniqueness Eigengroup 14 | -0.05 | -4.00 | 3.91 | 0.00 | 0.98 |
| Euler Number | 0.02 | 0.00 | 0.03 | 3.02 | 0.08 |
| Age, centred at 12 years | 1.32 | 0.82 | 1.81 | 27.25 | <.001*** |

*Note.* This model included Eigengroup uniqueness at previous timepoint (~4 - 44months before K10 timepoint). *$p$ <.05, **$p$≤.01, ***$p$≤.001

**Table S8: Lagged Wellbeing GEE by eigengroup uniqueness, age, sex, for *N*=116, *n*=497 datasets**

| Parameter | B | 95% CI | | Hypothesis Test | |
|---|---|---|---|---|---|
| | | Lower | Upper | Wald Chi-Square | Sig. |
| (Intercept) | 75.34 | 57.13 | 93.55 | 65.76 | <.001*** |
| [sex=Female] | 2.03 | -0.20 | 4.25 | 3.20 | 0.07 |
| Uniqueness Eigengroup 6 | 30.73 | 12.72 | 48.74 | 11.18 | <.001*** |
| Uniqueness Eigengroup 7 | -2.86 | -17.71 | 11.99 | 0.14 | 0.71 |
| Uniqueness Eigengroup 8 | -5.63 | -18.38 | 7.12 | 0.75 | 0.39 |
| Uniqueness Eigengroup 9 | -9.17 | -21.13 | 2.78 | 2.26 | 0.13 |
| Uniqueness Eigengroup 10 | 6.58 | -4.68 | 17.84 | 1.31 | 0.25 |
| Uniqueness Eigengroup 11 | -5.51 | -15.00 | 3.98 | 1.30 | 0.26 |
| Uniqueness Eigengroup 12 | 1.80 | -6.98 | 10.58 | 0.16 | 0.69 |
| Uniqueness Eigengroup 13 | 1.05 | -7.30 | 9.39 | 0.06 | 0.81 |
| Uniqueness Eigengroup 14 | 2.11 | -4.90 | 9.11 | 0.35 | 0.56 |
| Euler Number | -0.02 | -0.06 | 0.02 | 1.20 | 0.27 |
| Age, centred at 12 years | -1.61 | -2.45 | -0.78 | 14.33 | <.001*** |

*Note.* This model included Eigengroup uniqueness at previous timepoint (~4 - 44months before K10 timepoint). *$p$ <.05, **$p$≤.01, ***$p$≤.001

**Table S9: Mental health scale descriptives by age and sex, N=116**

| Timepoint | K10 mean (SD) | | COMPAS W mean (SD) | |
|---|---|---|---|---|
| | Female | Male | Female | Male |
| 1 | 17.22 (5.98) | 14.82 (4.32) | 101.16 (12.31) | 100.75 (10.52) |
| 2 | 15.43 (5.76) | 13.98 (3.24) | 102.11 (11.51) | 100.95 (10.73) |
| 3 | 16.33 (6.35) | 15 (4.38) | 100.9 (9.18) | 100.69 (10.02) |
| 4 | 16.82 (7.22) | 15.61 (5.83) | 101.71 (14) | 99.55 (9.97) |
| 5 | 17.11 (7.48) | 15.64 (6.28) | 100.11 (10.53) | 98.84 (12.33) |
| 6 | 15.43 (5.69) | 17.31 (9.83) | 100.71 (8.78) | 95.57 (15) |
| 7 | 20.11 (10.62) | 15.04 (6.39) | 97.79 (16.4) | 99.19 (12.16) |
| 8 | 17.67 (7.34) | 16.16 (6.01) | 99.78 (17.24) | 98.74 (14.39) |
| 9 | 22.43 (8.69) | 17.35 (6.54) | 94.14 (14.4) | 96.33 (17.44) |
| 10 | 19.85 (9.47) | 15.27 (4.63) | 98.31 (10.8) | 97.64 (14.55) |
| 11 | 23.93 (9.83) | 16.17 (5.31) | 96.53 (14.5) | 98.00 (11.86) |
| 12 | 23.79 (9.83) | 19.73 (7.14) | 96.00 (11.26) | 93.27 (18.59) |
| 13 | 21.31 (8.71) | 19.91 (9.92) | 97.00 (11.14) | 96.36 (12.45) |
| 14 | 16.62 (6.67) | 21.86 (8.28) | 103.77 (10.58) | 94.71 (14.12) |
| 15 | 20.50 (6.79) | 18.75 (11.53) | 98.50 (12.77) | 95.00 (19.44) |

*Note.* COMPAS-W includes $n$=610 datasets; K10 includes $n$=607 datasets
